# High Fat Diet-Induced Obesity Negatively Affects Whole Bone Bending Strength but not Cortical Structure in the Femur

**DOI:** 10.1101/729624

**Authors:** Nicholas J. Hanne, Andrew J. Steward, Jason M. Cox, Elizabeth D. Easter, Hannah L. Thornburg, Marci R. Sessions, Sriharsha V. Pinnamaraju, Jacqueline H. Cole

## Abstract

Although body mass index is positively associated with bone mineral density, suggesting obesity is protective against fracture, elderly obese individuals experience greater fracture risk at certain sites than non-obese peers, suggesting bone structural or material changes contribute to fragility. Diet-induced obesity rodent studies have reported detrimental changes to bone microstructure and some apparent-level material properties, but tissue-level material changes are not well understood. Because adipose tissue is highly vascularized, and bone remodeling depends critically on functional vascular supply, concurrent effects on osteovascular perfusion and structure may provide insight about obesity-related bone fragility. This study aimed to determine the effects of obesity on both tissue-level bone properties and osteovascular properties that could negatively impact bone strength. Five-week-old male C57Bl/6J mice were fed either high fat diet (HFD) or control fat diet (CFD) for 17 weeks and received daily treadmill exercise or remained sedentary for eight weeks at ages 14-22 weeks. HFD negatively affected femur bending strength, with 18% lower yield load than CFD. Although HFD negatively altered cancellous microstructure in the distal femur, with 32% lower bone volume fraction than CFD, it did not affect cortical bone geometry in the femoral metaphysis or diaphysis. HFD caused increased carbonate substitution but had no effect on other composition metrics or apparent- or tissue-level material properties in the femoral diaphysis. Exercise did not affect bone strength or microstructure but increased endosteal mineralizing surface in the tibial diaphysis, mineral crystallinity and mineral-to-matrix ratio in the femur, and blood supply to the proximal tibial metaphysis. HFD did not affect blood supply in the tibia or 2D osteovascular structure in the distal femoral metaphysis, indicating that HFD negatively affects cancellous bone without affecting osteovasculature. This study reveals that HFD negatively affected cancellous microstructure without affecting osteovascular structure, and whole-bone strength without altering cortical geometry or material properties.

## 1 Introduction

Over half of adults worldwide are overweight or obese.^(1)^ Higher bone mineral density (BMD), a primary determinant of bone strength^(2)^ that is associated with decreased fracture incidence in elderly men and women,^(3, 4)^ is associated with increasing body mass index (BMI) in both obese and non-obese individuals.^(4–7)^ However, increasing BMI in obese women is not as strongly correlated with increasing BMD and estimated material strength compared to non-obese women.^(7, 8)^ A meta-analysis reported that, despite having higher BMD, elderly obese individuals experience higher fracture incidence at particular sites compared to non-obese individuals – obese postmenopausal women have a higher risk of fracture in the spine, humerus, and leg bones but a lower risk of fracture in the hip and wrist, while older obese men have a higher risk of non-spinal fractures but a lower risk of fracture in the spine.^(5)^ Since bone strength depends not only on BMD but also structural and material properties,^(9, 10)^ the differential fracture risk with obesity likely results from adverse changes to bone structure and/or material properties, although these effects are understudied in humans. In non-obese elderly women, mid-tibial cortical thickness and cortical area, measured with high-resolution peripheral quantitative computed tomography (HR-pQCT), were positively correlated with BMI.^(11)^ Despite these beneficial changes to geometry, cortical bone material strength index (BMSi) in the tibia, measured with reference point indentation, had a weak negative correlation with both BMI and subcutaneous fat in the tibia.^(11)^ Examining bone structure and material properties beyond HR-pQCT and BMSi is difficult in humans, but they have been examined in animal models of obesity. Previous diet-induced obesity studies in young, mostly male, mice reported detrimental changes to trabecular microstructure in the femur,^(12–17)^ cancellous bone formation rate,^(17)^ serum concentrations of osteocalcin, tartrate resistant acid phosphatase,^(14)^ and carboxyl-terminal collagen crosslinks,^(12)^ and cortical apparent material properties (bending apparent modulus and ultimate stress) and fracture toughness,^(18, 19)^ but no change to femoral areal BMD,^(18, 19)^ cortical volumetric BMD (vBMD),^(16)^ or cancellous tissue mineral density (TMD)^(14–16)^ for high fat diet (HFD) compared to control fat diet (CFD). Therefore, HFD-induced obesity induces some structural and apparent-level material changes without changes to bone density, further supporting the notion of tissue-level effects that need to be further examined.

Vascular properties may contribute to the detrimental changes in cancellous bone structure with obesity. Adipose and bone tissues are highly vascularized and require adequate blood flow for formation and homeostasis.^(20–23)^ In rodent studies, HFD increases the amount of adipose in the medullary cavity of long bones,^(13,24–27)^ and adipose produces angiogenic cytokines that induce rapid vascularization.^(20, 28)^ Although increased bone vascularization is associated with increased bone formation rate in cancellous bone in normal-weight rats,^(29)^ HFD is associated with detrimental changes to cancellous bone structure.^(12–17)^ In addition to the amount of blood vessels, the structure of vasculature within bone is also important for remodeling; compared to non-remodeling bone surfaces, active sites of bone remodeling have increased number of capillaries within 50 µm of the bone surfaces.^(30, 31)^ Exercise reduces the accumulation of adipose within the long bones of mice fed HFD^(24, 25)^ and stimulates osteovascular crosstalk pathways, such as VEGF and bone morphogenetic protein 2 (BMP2), that promote bone formation.^(32, 33)^ However, the effects of HFD and exercise on the osteovasculature is understudied. In this study, we examined changes in both bone and osteovascular tissues using a mouse model of diet-induced obesity, both with and without moderate treadmill activity. We hypothesized that obesity decreases the integrity of bone microstructure and material properties, while exercise induces new vascular and bone growth.

## 2 Materials and Methods

### 2.1 Study Design

The protocol for this work was approved by the Institutional Animal Care and Use Committee at North Carolina State University. Sixteen 5-week-old male C57Bl/6J mice (The Jackson Laboratory, Bar Harbor, ME) were fed a high fat diet (D12492 60% kcal fat, Research Diets, Inc., New Brunswick, NJ) (n=8, “HFD” group) or a matched control fat diet (D12450B 10% kcal fat, Research Diets, Inc) (n=8, “CFD” group) for 17 weeks (Figure 1). Mice were housed with their groups (4-5 per cage) under controlled 12-hour diurnal photoperiod and fed their respective diets *ad libitum*. After 9 weeks of diet (Week 9, 14 weeks of age), after the obesity phenotype was established, mice were further divided into two activity groups, either daily treadmill exercise (n=4 from each diet group, “CFD-Exercise” and “HFD-Exercise”) or stationary treadmill groups (n=4 from each diet group, “CFD-Sedentary” and “HFD-Sedentary”).

**Figure 1:**
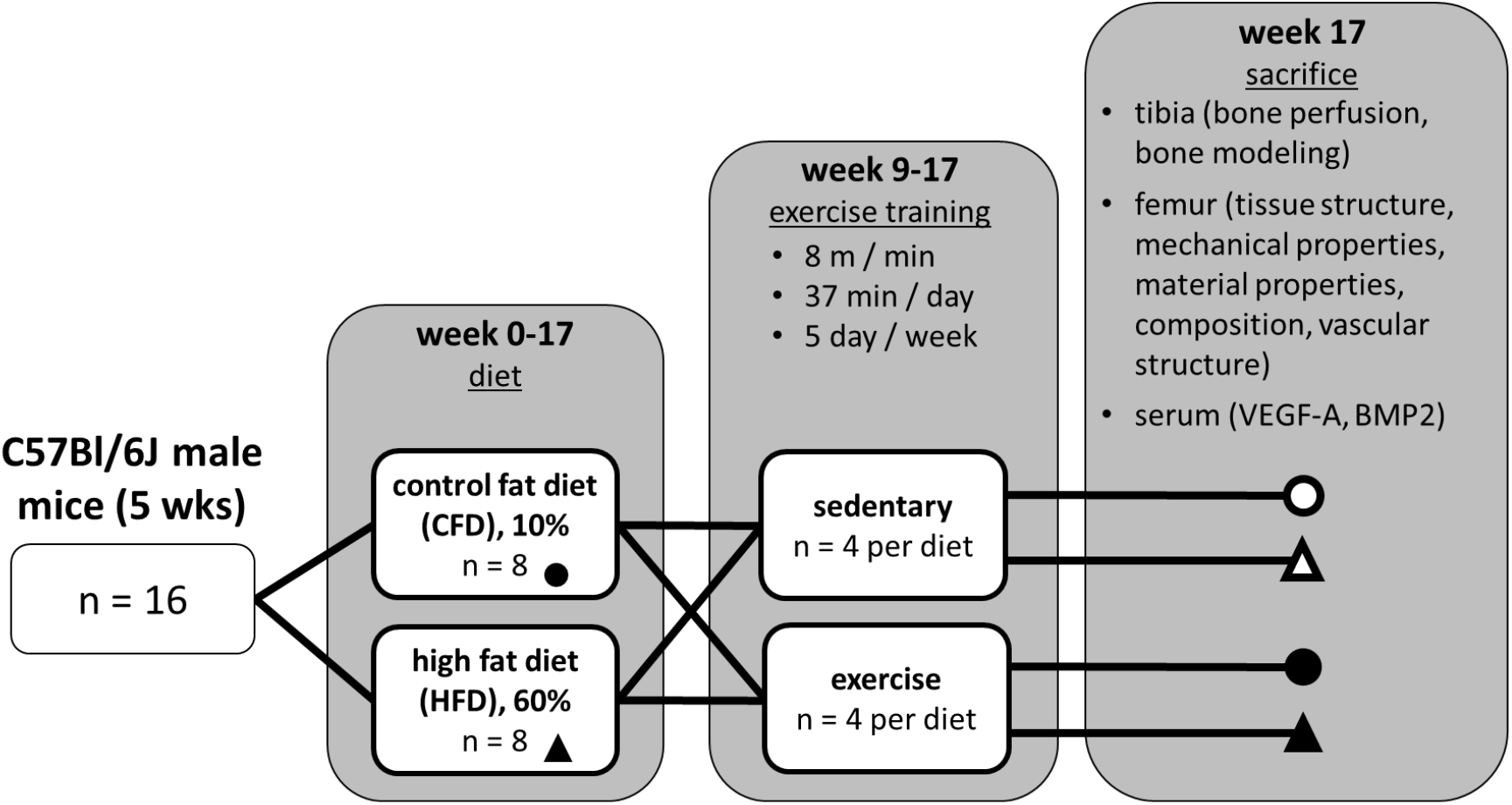
Experimental design: Mice were fed either control fat diet (CFD) or high fat diet (HFD) starting at 5 weeks of age. After 9 weeks of diet, groups either were exercised (moving treadmill) or remained sedentary (stationary treadmill) for 8 weeks. After 17 total weeks of diet and 8 weeks of exercise, various endpoint vascular and bone metrics were analyzed.

Exercise mice were acclimated to a mouse treadmill (Exer 3/6, Columbus Instruments, Columbus, OH) over three days of increasing speeds (day 1: 6 m/min for 10 min, day 2: 9 m/min for 10 min, day 3: 12 m/min for 10 min). After acclimation, exercise groups ran on the treadmill 5 days per week for 8 weeks (8 m/min for 37 min at a 5-degree incline). Mice in the HFD group were unable to run for 30 min at 10 m/min, so the protocol was adjusted to 8 m/min for a longer time to provide the same running distance (300 m). Sedentary groups were placed on an immobile replica treadmill for the same duration as the exercise groups. Exercise and diet were continued for 8 weeks until the end of the study (Week 17, 22 weeks of age).

For dynamic histomorphometry, alizarin complexone (C0875, Sigma-Aldrich, St. Louis, MO) and calcein (A3882, Sigma-Aldrich) were injected intraperitoneally (30 mg/kg) at 10 and 3 days prior to sacrifice, respectively. At the conclusion of the study, and immediately before sacrifice, *in vivo* measurements of tibial perfusion were made under anesthesia (described below). Mice were euthanized by CO2 asphyxiation followed by cervical dislocation. For serum assays, blood was immediately collected through cardiac puncture and left at room temperature for 30 min to clot, after which the serum was separated by centrifugation (2,000 x g for 10 min) and stored at −80°C. The left and right femora and tibiae were dissected. The left femur and both tibiae were fixed in 10% neutral buffered formalin at 4°C for 36 hours, then stored in 70% ethanol at 4°C. The unfixed right femur was wrapped in 1X phosphate buffered saline (PBS)-soaked gauze and fresh frozen at −20°C.

### 2.2 Obesity Phenotype

Body mass and serum glucose were measured weekly in all groups following the initiation of treadmill exercise in Week 9. Serum glucose concentration was measured from the tail vein after 6 hours of fasting (AlphaTrak 2 Blood Glucose Monitoring System, Abbott Laboratories, Abbott Park, IL). Glucose tolerance tests (GTT) were performed following 6 hours of fasting at Week 13 and Week 17 to assess ability to clear a bolus injection of glucose from the blood, which is a test for the development of diabetes. For the test, a 0.3 g/mL (30%) glucose solution was injected intraperitoneally at 1 g of glucose per kg of body mass. Serum glucose concentration was measured immediately prior to the injection of glucose and 15, 30, 60, 90, and 120 min after the injection. Glucose concentrations over the maximum threshold of the glucometer were recorded as 750 mg/dL (upper range concentration of the AlphaTrak 2). The areas under the curve for the GTT results were calculated using the trapezoid rule.

### 2.3 Bone Perfusion (Tibia)

*In vivo* tibial perfusion was measured at the endpoint of the study in the right proximal tibial metaphysis with laser Doppler flowmetry (LDF). LDF can quantify vascular perfusion – a functional measure of bone blood flow comprised of amount of vasculature, velocity and direction of blood flow, and vascular permeability – in murine tibiae.^(34, 35)^ Perfusion readings were taken just prior to sacrifice using an LDF monitor with 785-nm light source and selectable 3 kHz lowpass filter (moorVMS-LDF, Moor Instruments Ltd., Axminster, UK) paired with a needle probe (VP4, 0.8 mm outer diameter, 0.25 mm fiber separation), as follows. After 6 hours of fasting, mice were anesthetized with 2% isoflurane in pure oxygen. Mice were placed supine, and the shaved right hindlimb was taped to a heated surgical platform. A 2-5 mm long was made over the anteromedial side of the proximal tibial metaphysis, the periosteum was gently scraped away from the metaphysis, and the LDF probe was held flush to the bone with a micromanipulator (MM3-ALL, World Precision Instruments, Sarasota, FL) for a 30-second recording. The probe was removed and replaced two more times, and the weighted mean of the three recordings was used for analysis. Measurements are expressed in perfusion units (PU), arbitrary units that are standard for LDF.

### 2.4 Cancellous and Cortical Bone Structure (Femur)

Cancellous bone microstructure and cortical bone geometry were assessed in the left femur by scanning in 70% ethanol with micro-computed tomography (micro-CT, µCT80, SCANCO Medical AG, Brüttisellen, Switzerland) using a 10-µm voxel size, 45 kV peak X-ray tube potential, 177 µA X-ray intensity, and 800-ms integration time. Volumes of interest (VOI) were analyzed using the scanner’s software (SCANCO v.6.6) for standard cortical and cancellous bone metrics.^(36)^ The distal metaphyseal VOI was defined as 10% of the total femur length positioned proximal to the distal growth plate. The cancellous and cortical bone were contoured and analyzed separately in the metaphysis. The diaphyseal VOI was defined as 15% of the total femur length, centered between the distal growth plate and the middle of the third trochanter. The mid-diaphyseal VOI (used for estimating apparent-level material properties with three-point bending data) was defined as a 2.5-mm section with the same center as the diaphyseal VOI.

### 2.5 Cortical Bone Remodeling (Tibia)

Dynamic indices of cortical bone remodeling were examined in the right tibial diaphysis using dynamic histomorphometry. The right tibia was embedded in methylmethacrylate, then sectioned transversely in 200-µm thick sections just distal to the tibiofibular junction under constant water irrigation using a low-speed precision saw (IsoMet Low Speed Precision Cutter, Buehler, Lake Bluff, IL). Sections were glued to glass slides with cyanoacrylate glue and sanded to 10-30 µm thickness with increasing grit sandpaper.^(37)^ Two sections from each bone were imaged at 40X on a Zeiss LSM 880 laser scanning microscope with Airyscan (Carl Zeiss Microscopy, Thornwood, NY). Standard dynamic histomorphometry parameters – mineralizing surface per bone surface (MS/BS), mineral apposition rate (MAR), and bone formation rate (BFR/BS) – were measured on two sections per mouse using ImageJ (version 1.51v) and Photoshop (version CC 2018m, Adobe Systems Inc., San Jose, CA),^(38, 39)^ and the mean values were used for analysis.

### 2.6 Whole Bone and Apparent-Level Mechanical Properties (Femur)

The right femur underwent three-point bending to failure to measure whole bone mechanical properties and estimated apparent-level material properties. Immediately prior to testing, the femur was brought to room temperature and placed in a 37°C bath of 1X PBS for 60 sec. The bone was centered over a 6.5-mm lower span (40% average femur length) with the anterior side facing up so that the anterior diaphysis was loaded in compression. Three-point bending was performed to failure using an actuator speed of 0.025 mm/sec (EnduraTec ELF 3220, Bose Corp., Minnetonka, MN). Force (500-g capacity load cell, Sensotec Model 31/6775-06, Honeywell Sensotec, Columbus, OH) and displacement were recorded at 100 Hz. After failure, the femur was immediately wrapped in PBS-soaked gauze and returned to −20°C. Yield load (F_yield_), maximum (ultimate) load (F_ultimate_), stiffness, post-yield deformation (PYD), and work-to-fracture were calculated from load-displacement curves with MATLAB^®^ (R2017, The MathWorks, Inc., Natick, MA).^(40)^ Yield was calculated as the point where a line with a 5% decrease in stiffness intersected the force-displacement curve.^(40)^ PYD was calculated as the difference between the deformation at yield and the deformation at failure. The stress-strain curve was estimated using the cross-sectional moment of inertia about the bending axis calculated from the mid-diaphyseal VOI in the micro-CT scans of the left femur (described above).^(41)^ Yield stress (σ_yield_), maximum (ultimate) stress (σ_ultimate_), and Young’s modulus (E) were calculated from these estimated stress-strain curves with MATLAB^®^.

### 2.7 Tissue-Level Mechanical Properties (Femur)

Cortical bone material properties were examined with nanoindentation in the right femoral diaphysis. Following three-point bending, the right femur was already divided in half at the failure point (position of the top loading point); a 1-2 mm transverse section was cut just distal to the distal half and affixed to a glass slide with the fractured end facing up. The remainder of the distal half of the right femur was reserved for immunofluorescence (described below). The section was smoothed with increasing grit sandpaper (120 followed by 600 grit) and then polished with 3 µm diamond slurry (90-3DL3, Allied High Tech Products Inc, Rancho Dominguez, CA) until smooth.^(42)^ Before nanoindentation, Raman spectroscopy was performed on the proximal surface of the polished cortical section (described below). Then nanoindentation was performed on the same samples using a Hysitron TriboIndenter TI 980 with a diamond Berkovich tip (Bruker, Billerica, MA). The instrument was calibrated by performing indentations in air and a fused quartz standard. Each bone was indented in the anterior and posterior regions of the mid-cortex in a 4×4 grid of points equally spaced 15 µm apart. A trapezoidal loading function with 60 sec loading to 3000 µN, 30 sec holding at peak load, and 6 sec unloading was performed at each point.^(42, 43)^ The fused quartz standard was tested before and after each mid-cortex grid to validate calibration and remove organic material from the tip. Force-displacement curves exhibiting nonlinearity during loading were removed from analysis. Hardness (H) and reduced modulus (E_r_) were calculated from the force-displacement curve during unloading and were averaged across each region grid, giving a mean for each anterior and posterior region.^(44)^

### 2.8 Cortical Bone Composition (Femur)

Cortical bone tissue composition was measured with Raman spectroscopy (XploRA PLUS confocal Raman microscope, HORIBA Scientific, Piscataway, NJ). Raman spectra were collected with a 785-nm laser at 50X magnification at the endosteal edge, mid-cortex, and periosteal edge in the posterior, lateral, anterior, and medial quadrants of the section (Figure 2A). Mid-cortex quadrant scans were comprised of a 2 x 5 grid of point collections spaced 5 µm apart, while endosteal and periosteal quadrant scans were comprised of a line of 6 points spaced 2 µm apart, aligned parallel to and positioned 5-10 µm in from the bone surface. Each point was a 30-second accumulation in the 800-1800 cm^-1^ range. The spectrometer software (LabSpec 6, v.6.5.1.24) automatically performed baseline correction, while the remaining analysis was performed in MATLAB^®^.

**Figure 2:**
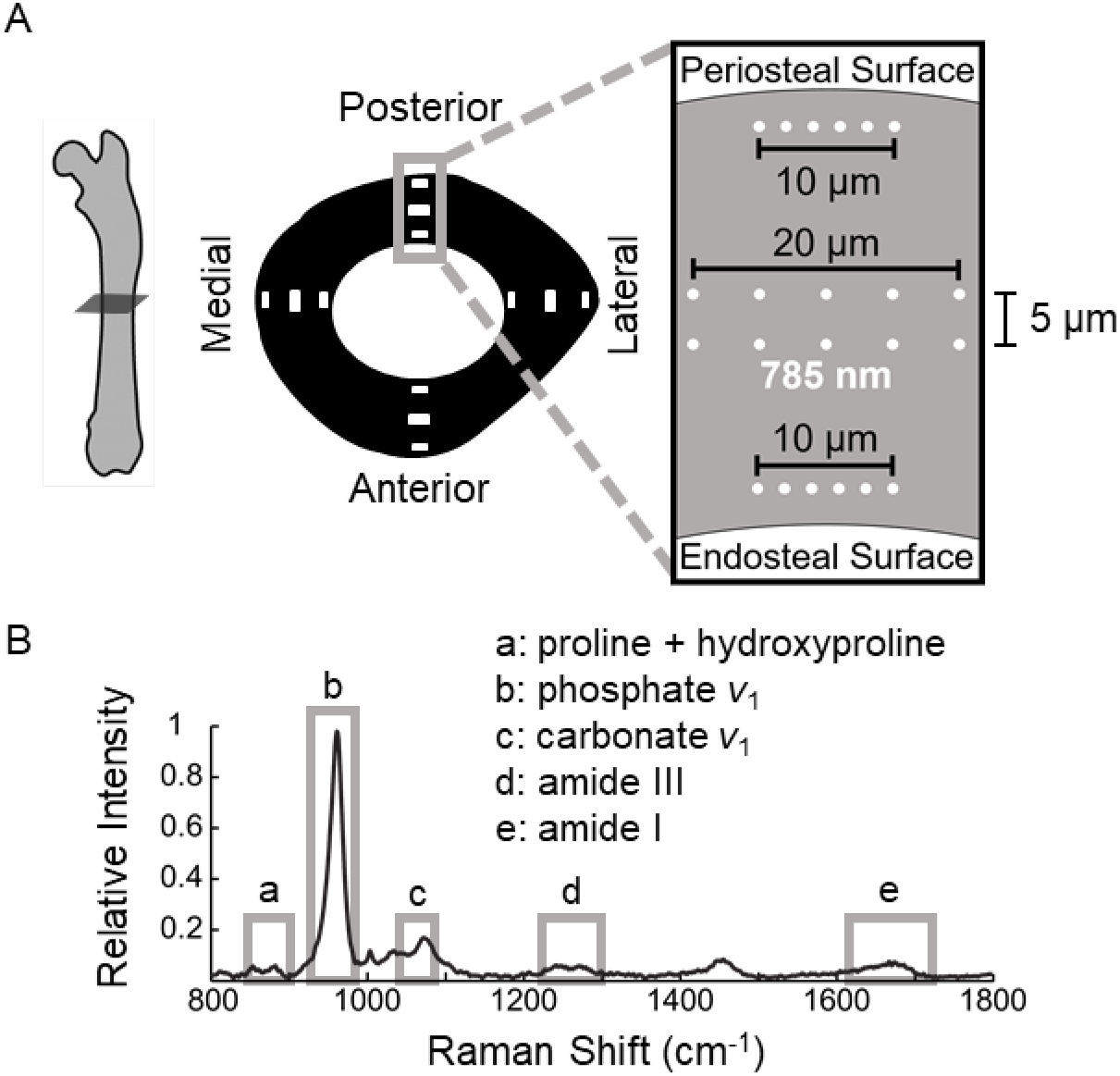
A) Bone composition was assessed by Raman spectroscopy at three positions in the posterior, lateral, anterior, and medial quadrants within the cortical diaphysis of right femora: mid-cortex (2×5 point array) and endosteal and periosteal edges (1×6 linear array). B) Raman spectra were normalized to the phosphate *v*_1_ band intensity (b), and crystallinity was calculated as the inverse of the full-width at half maximum of the phosphate *v*_1_ band. Mineral-to-matrix band intensity ratios were calculated for phosphate *v*_1_ relative to the summation of proline and hydroxyproline (a), amide I (e), and amide III (d). Carbonate substitution (carbonate *v*_1_ (c)/ phosphate *v*_1_) and carbonate-to-matrix ratio (carbonate *v*_1_ / amide I) were also calculated.

Spectra were normalized relative to the phosphate *v*_1_ maximum intensity (930-980 cm^-1^), and the maximum normalized intensities were determined in the regions corresponding to the summation of proline (830-863 cm^-1^) and hydroxyproline (864-899 cm^-1^), phosphate *v*_1_ (930-980 cm^-1^), carbonate *v*_1_ (1055-1090 cm^-1^), amide III (1220-1300 cm^-1^), and amide I (1616-1720 cm^-1^) (Figure 2B).^(45, 46)^ Several standard metrics were calculated, as follows.^(45)^ Mineral-to-matrix ratios were calculated as the ratio of the phosphate *v*_1_ normalized intensity relative to amide I, amide III, or summed proline and hydroxyproline normalized intensity. The carbonate-to-matrix ratio was calculated as the ratio of the carbonate *v*_1_ to amide I normalized intensities. Carbonate substitution was calculated as the normalized intensity of carbonate *v*_1_. Mineral maturity (crystallinity) was calculated as the inverse of the full-width at half maximum (FWHM) of a single-order Gaussian curve fit to the phosphate *v*_1_ band. Each of these Raman metrics were averaged across each measurement grid within quadrants, giving a mean for each region (endosteal edge, mid-cortex, or periosteal edge) at each quadrant (posterior, lateral, anterior, and medial).

### 2.9 Osteovascular Structure (Femur)

Vascular structure and proximity to bone surfaces were examined in the distal femoral metaphysis using thick-section immunofluorescence to quantify the amount of blood vessels, labeled by endomucin (EMCN), and bone surfaces, labeled by collagen type I (COL-1).^(47)^ The remaining distal portion of the right femur samples were fixed overnight in 10% neutral buffered formalin at 4°C, decalcified in 0.5M ethylenediaminetetraacetic acid at 4°C for 24 hours, and then embedded in cryoprotectant embedding media comprised of 8% gelatin (G1890, Sigma-Aldrich), 2% polyvinylpyrrolidone (P5288, Sigma-Aldrich), and 20% sucrose (S7903, Sigma-Aldrich) in 1X PBS. Samples were sectioned longitudinally in 100-µm thick sections on a cryotome at −23°C (HN 525NX, Thermo Fisher Scientific, Waltham, MA). Sections were stained overnight at 4°C using unconjugated primary antibodies at 1:100 dilution for endomucin (rat anti-mouse sc-65495, Santa Cruz Biotechnology, Santa Cruz, CA) and at 1:200 dilution for collagen type I (rabbit anti-mouse, AB765P, MilliporeSigma, Burlington, MA). Secondary antibodies at 1:200 dilution were added for 90 min at room temperature (goat anti-rat with AlexaFluor 647 ab150159, Abcam, Cambridge, UK; goat anti-rabbit with AlexaFluor 488 A11006, Invitrogen, Carlsbad, CA). DAPI at 2 µg/mL was added for 10 min at room temperature to counterstain nuclei. All sections were imaged at 20X on a Zeiss LSM 880 laser scanning microscope with Airyscan. Regions with positive COL-1 and EMCN labeling were traced by hand in ImageJ (version 1.51v) in a region of interest (ROI) that was 10% of the femur length and positioned just proximal to the distal growth plate (same as the micro-CT metaphyseal VOI). Vascular structure was analyzed by calculating EMCN^+^ area per total area, COL-1^+^ area per total area, and the distance between EMCN^+^ and COL-1^+^ surfaces in MATLAB^®^. Several samples were destroyed or lost during sectioning, so only a subset of samples were analyzed (n = 1 CFD-Sedentary, n = 3 CFD-Exercise, n = 2 HFD-Sedentary, n = 1 HFD-Exercise).

### 2.10 Osteovascular Crosstalk (Serum)

To examine osteovascular crosstalk between endothelial cells and osteoblasts, serum concentrations of bone morphogenetic protein 2 (BMP2) and vascular endothelial growth factor A (VEGF-A) were measured using serum collected and stored at the endpoint of the study. Serum concentrations were measured with enzyme-linked immunosorbent assays (ELISA) per the manufacturers’ instructions, using mouse-specific kits for BMP2 (ab119582, Abcam) and VEGF-A (KMG0111, Thermo Fisher Scientific). All samples were analyzed in duplicate using a plate reader (Synergy H1M, BioTek Instruments, Inc., Winooski, VT).

### 2.11 Statistical Analyses

All statistical models were analyzed in SAS (SAS University Edition v. 9.4, SAS Institute Inc., Cary, NC) or R (R v. 3.5.1, R Foundation for Statistical Computing, Vienna, Austria) to determine the following: 1) effects of HFD and exercise on body mass and fasting serum glucose concentration at each week after treadmill exercise was started; 2) effects of HFD and exercise on metrics of glucose tolerance, bone perfusion, cancellous and cortical bone microstructure, cortical bone remodeling, whole bone mechanical properties, apparent-level material properties, osteovascular structure, and osteovascular crosstalk; 3) effects of HFD and exercise on cortical bone material properties measured with nanoindentation and composition measured with Raman spectroscopy. All data are presented as the group mean ± standard deviation unless otherwise stated. Results from nanoindentation and Raman spectroscopy are presented as mean across the scanned regions.

For analysis #1, mouse mass and serum glucose were compared between diet and activity groups across weekly timepoints using a repeated measures factorial model with interaction between all terms (SAS ‘MIXED’ procedure). Diet (CFD or HFD) and activity (sedentary or exercise) were modeled as fixed factors, while week was modeled as a repeated measure within each mouse. The residual variance was modeled assuming compound symmetry covariance, chosen as the covariance structure that provided the best fit to the data. Predicted least-squares means with Tukey-Kramer adjustment for multiple comparisons were used to analyze effect differences between diet and activity groups, with interaction, at each timepoint (i.e., CFD-Sedentary vs. HFD-Sedentary at Week 9).

For analysis #2, outcome parameters were compared between diet and activity, with interaction, using two-way analysis of variance (R ‘aov’ function). Tukey’s post-hoc tests were used to compare group means. Vascular structure parameters were analyzed with a similar model, but the interaction between diet and activity was not modeled due to missing data and thus insufficient power to analyze the full model. Three-point bending parameters were further analyzed with two analysis of covariance (ANCOVA) models, one with mass as the continuous covariate and one with femur length as the continuous covariate.^(40, 48)^

For analysis #3, the same repeated measures factorial model used in analysis #1 was used (SAS ‘MIXED’ procedure), but parameters were compared between diet and activity groups across scan region (anterior and posterior for nanoindentation; posterior, lateral, anterior, and medial for Raman spectroscopy). The residual variance was modeled assuming compound symmetry covariance. Predicted least-squares means with Tukey-Kramer adjustment for multiple comparisons were used to analyze pairwise differences between diet and activity groups, with interaction (i.e., HFD-Sedentary vs. HFD-Exercise).

## 3 Results

### 3.1 Obesity Phenotype

Weekly measures of mouse mass, serum glucose, and monthly glucose tolerance tests confirmed that the high fat diet produced an obese phenotype in this study. The HFD group had consistently greater body mass at all timepoints compared to the CFD group (p = 0.0016, Figure 3A). At the end of the study, after 17 weeks of diet, the HFD group (43.0 ± 5.2 g) weighed 33% more than the CFD group (32.4 ± 1.7 g, p < 0.0001). Overall, the HFD group had increased fasting glucose concentrations relative to the CFD group (p = 0.0054), but not at every timepoint (Figure 3B). At the end of the study, fasting glucose concentration was 27% higher in the HFD group (246 ± 28 mg/dL) than in the CFD group (193 ± 36 mg/dL, p = 0.014). Exercise did not affect body mass (p = 0.76) or fasting serum glucose concentration (p = 0.57).

**Figure 3:**
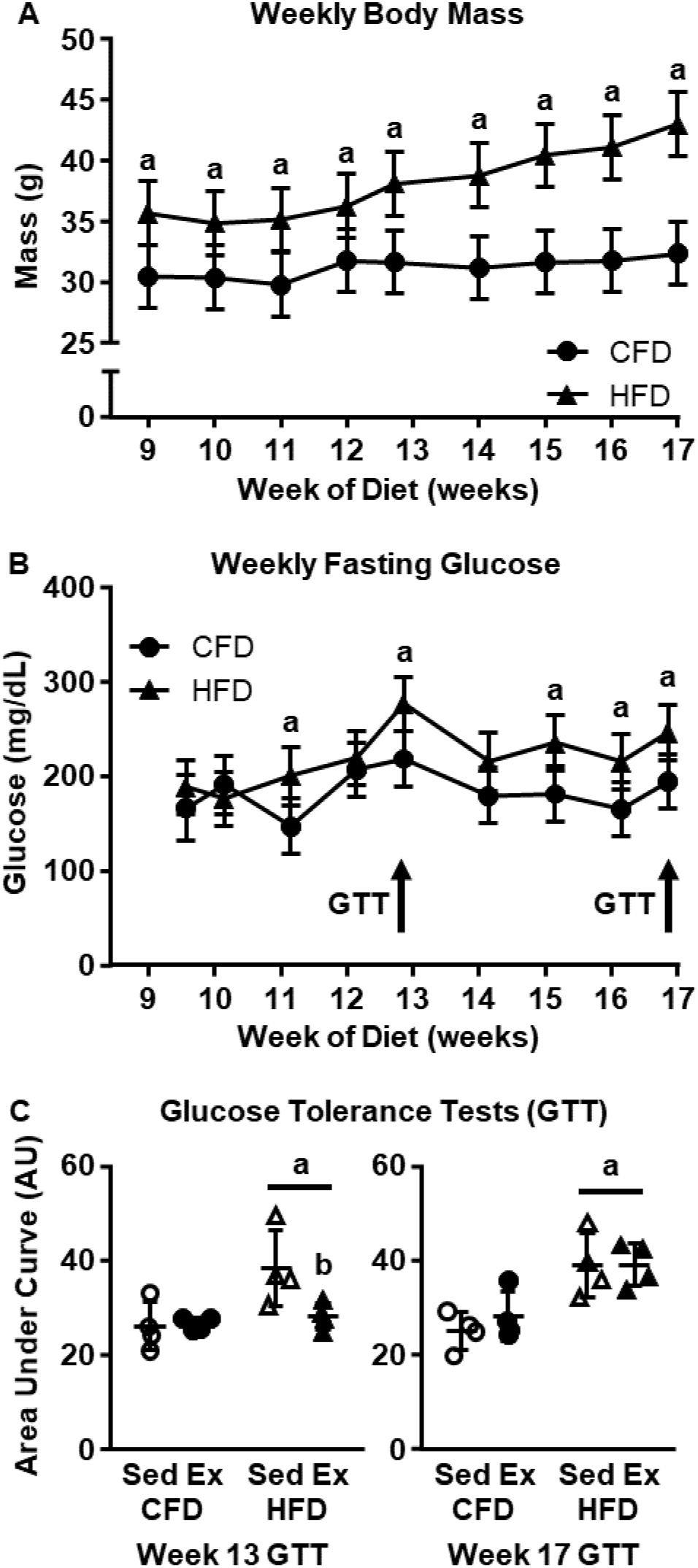
A) Body mass was consistently higher with HFD than CFD at every timepoint. B) Weekly fasting glucose concentration were higher in the HFD group at Week 11, 13, 15, 16, and 17. C) HFD had lower glucose tolerance (higher area under curve) than CFD at Weeks 13 and 17 of diet. Data in A-B presented as estimated least-squares mean ± 95% confidence interval. a: p < 0.05 HFD vs. CFD (main effect), b: p < 0.10 HFD-Exercise (Ex) vs. HFD-Sedentary (Sed).

The HFD group had a lower glucose tolerance, metabolizing a bolus of glucose more slowly (represented by larger area under the curve) than did the CFD group at Week 13 (HFD: 33.2 ± 9.0 p = 0.017 vs. CFD: 26.2 ± 2.2) and Week 17 (HFD: 39.0 ± 9.0, p = 0.0004 vs. CFD: 26.5 ± 7.4) (Figure 3C). At Week 13, exercise nearly improved glucose tolerance in the HFD-Exercise group (28.2 ± 2.8) relative to the HFD-Sedentary group (38.2 ± 8.1, p = 0.066), bringing the glucose tolerance of HFD-Exercise similar to that of CFD-Sedentary (26.0 ± 5.1, p = 0.92) and CFD-Exercise (26.0 ± 1.7, p = 0.96) groups. The benefit of exercise in the HFD group did not persist, however, and at Week 17, the HFD-Exercise (39.1 ± 4.5) and HFD-Sedentary (38.9 ± 6.7) groups had similar glucose tolerance (p = 1.00) elevated above that of the CFD groups (CFD-Exercise: 28.0 ± 5.2; CFD-Sedentary: 25.0 ± 3.9). Several mice had glucose concentrations that were above the detectible range of the glucometer, which artificially decreased the area under the curves.

During the Week 13 GTT, one HFD-Sedentary mouse had over-range readings at three timepoints, and one CFD-Sedentary mouse had over-range readings at one timepoint. During the Week 17 GTT, one HFD-Sedentary and one HFD-Exercise mouse had over-range readings at two timepoints each, and one CFD-Exercise had an over-range reading at one timepoint.

### 3.2 Bone Perfusion (Tibia)

At the end of the study, *in vivo* perfusion in the proximal tibial metaphysis was 29% greater in exercise groups (12.2 ± 3.0 PU) compared to sedentary groups (9.5 ± 1.6 PU, p = 0.044, Figure 4). Tibial perfusion was similar between HFD (11.7 ± 2.6 PU) and CFD groups (10.1 ± 2.7 PU, p = 0.23).

**Figure 4:**
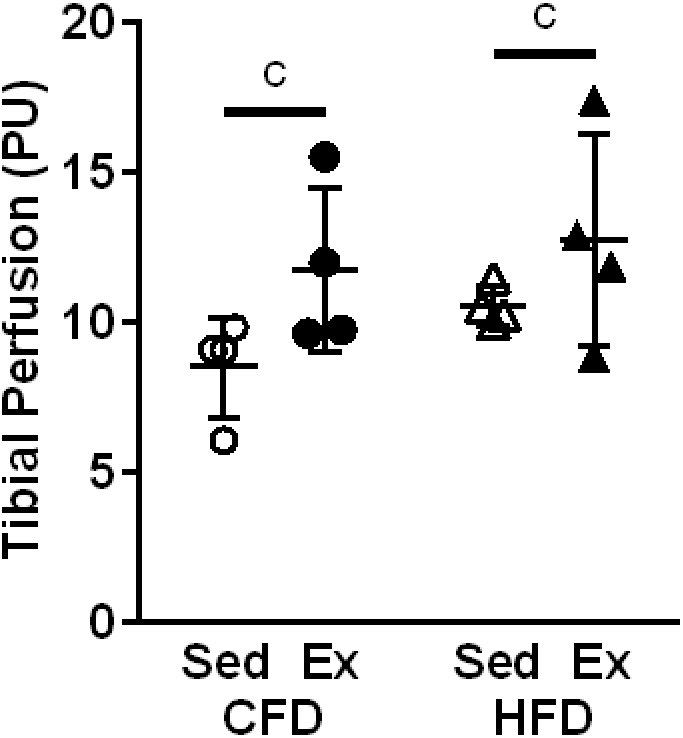
Bone perfusion in the proximal tibial metaphysis was significantly higher in exercise (Ex) than in sedentary (Sed) groups but not with HFD compared to CFD. c: p < 0.05 Ex vs. Sed (main effect). PU = perfusion unit (arbitrary).

### 3.3 Cancellous and Cortical Bone Structure (Femur)

High fat diet had detrimental effects on trabecular microstructure in the distal femoral metaphysis. Compared to CFD, the HFD group had 32% lower bone volume fraction (BV/TV, HFD: 11.2 ± 3.5% vs. CFD: 16.4 ± 3.2%, p = 0.0089, Figure 5A); 20% lower trabecular number (Tb.N, HFD: 3.73 ± 0.23 mm^-1^ vs. CFD: 4.64 ± 0.54 mm^-1^, p = 0.0010, Figure 5B); and 26% greater trabecular separation (Tb.Sp, HFD: 262.4 ± 18.4 µm vs. CFD: 208.5 ± 21.0 µm, p=0.0001, Figure 5C); but similar trabecular thickness (Tb.Th, HFD: 51.3 ± 6.6 µm vs. CFD: 50.1 ± 3.6 µm, p = 0.68, Figure 5D). In addition, the connectivity density (Conn.D) of the trabecular network was 50% lower in the HFD group compared to the CFD group (p = 0.00053, Table 1), but the degree of anisotropy (DA) was not significantly different between HFD and CFD groups (p = 0.11, Table 1). HFD group had a 30% lower trabecular vBMD than CFD group (p = 0.0082) but similar TMD (p = 0.98). Exercise did not significantly affect any of the metrics for trabecular bone microstructure.

**Figure 5:**
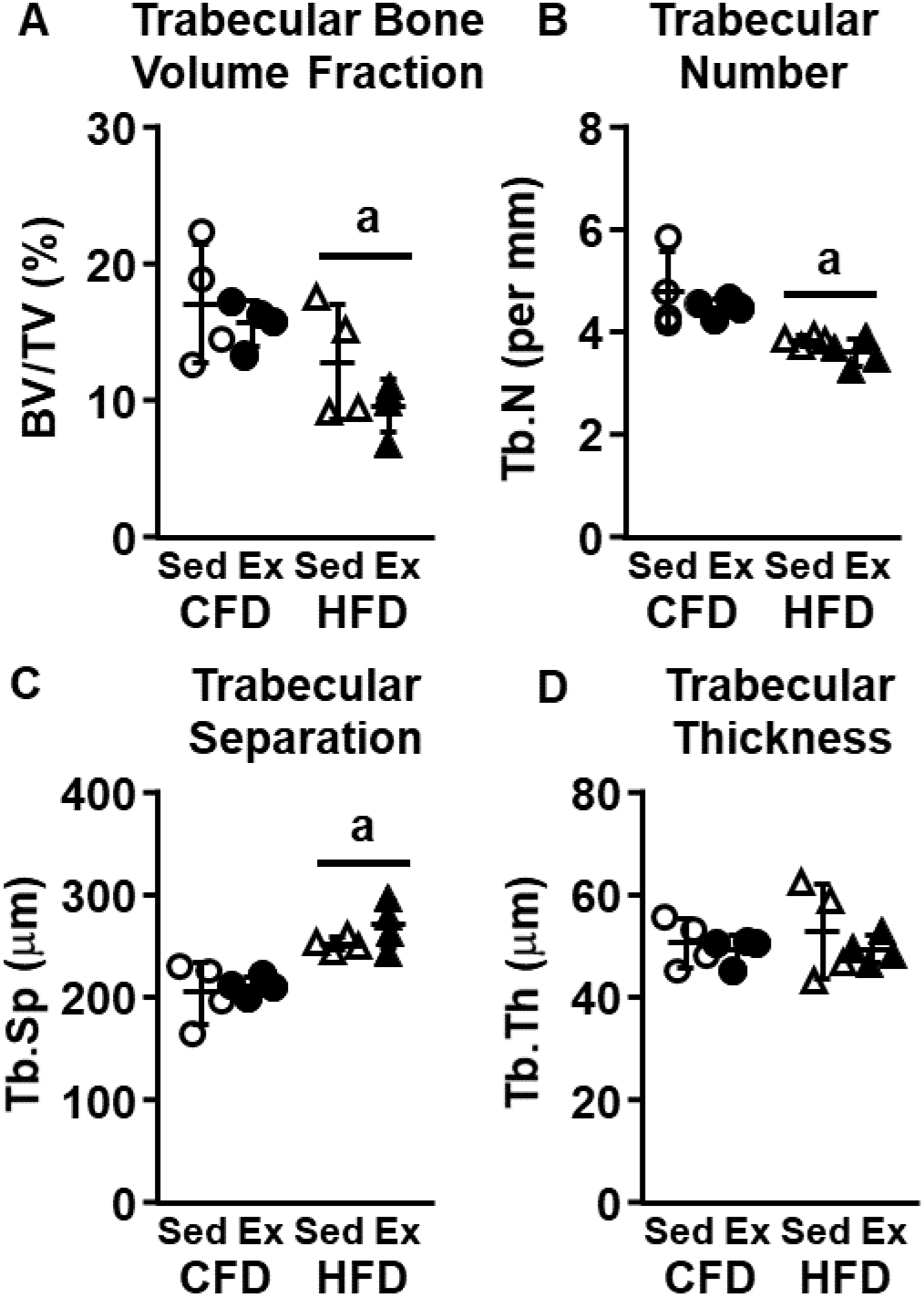
Relative to CFD, HFD groups exhibited significantly less robust trabecular architecture in the distal femur, with A) decreased bone volume fraction and B) trabecular number and C) increased trabecular separation, but D) no differences in trabecular thickness. a: p < 0.05 HFD vs. CFD (main effect).

**Table 1:**
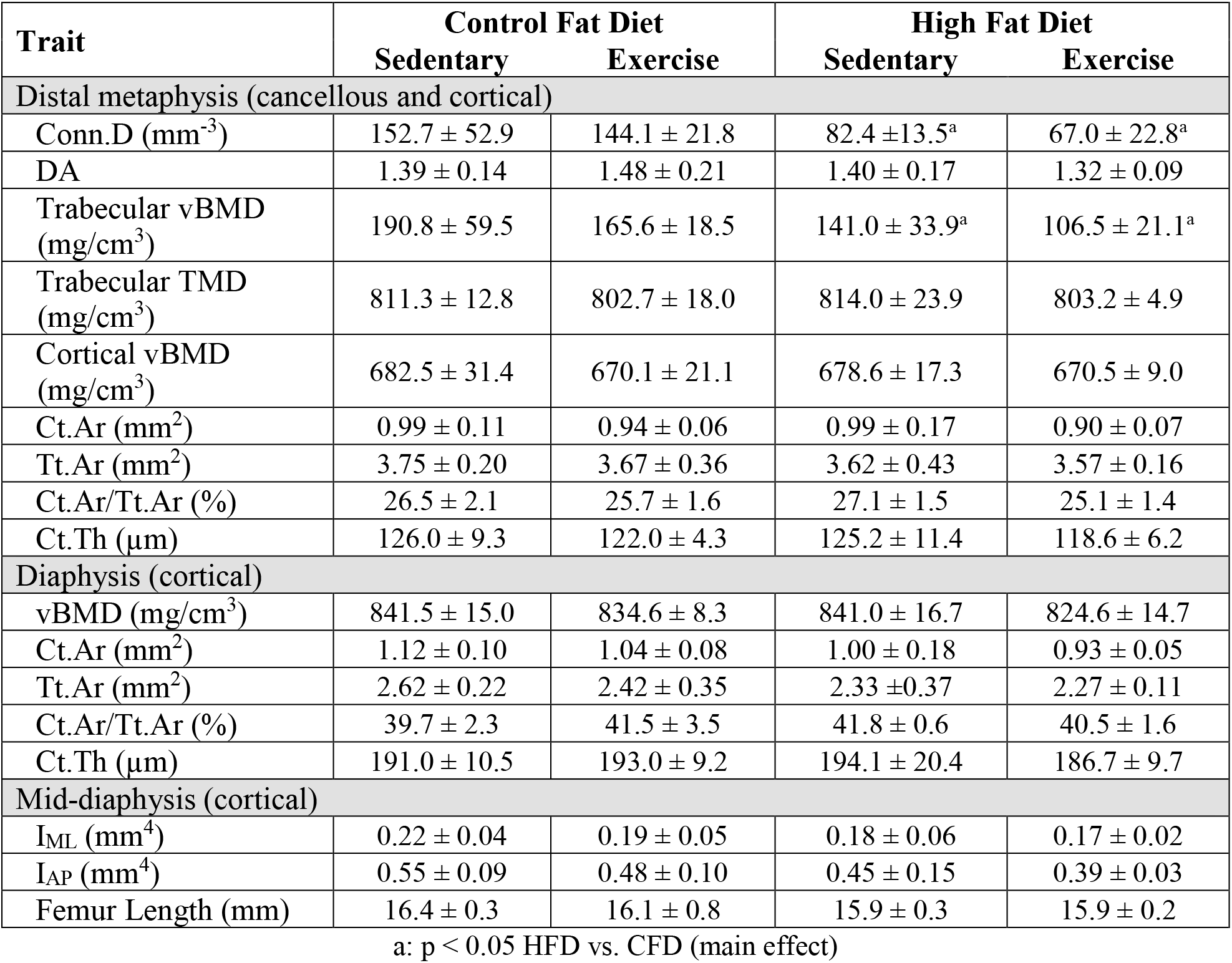
Cancellous and Cortical Bone Structure in the Femur (mean ± SD)

| Trait | Control Fat Diet |  | High Fat Diet |  |
| --- | --- | --- | --- | --- |
|  | Sedentary | Exercise | Sedentary | Exercise |
| Distal metaphysis (cancellous and cortical) |  |  |  |  |
| Conn.D ($\text{mm}^{-3}$) | 152.7 ± 52.9 | 144.1 ± 21.8 | 82.4 ± 13.5<sup>a</sup> | 67.0 ± 22.8<sup>a</sup> |
| DA | 1.39 ± 0.14 | 1.48 ± 0.21 | 1.40 ± 0.17 | 1.32 ± 0.09 |
| Trabecular vBMD ($\text{mg}/\text{cm}^3$) | 190.8 ± 59.5 | 165.6 ± 18.5 | 141.0 ± 33.9<sup>a</sup> | 106.5 ± 21.1<sup>a</sup> |
| Trabecular TMD ($\text{mg}/\text{cm}^3$) | 811.3 ± 12.8 | 802.7 ± 18.0 | 814.0 ± 23.9 | 803.2 ± 4.9 |
| Cortical vBMD ($\text{mg}/\text{cm}^3$) | 682.5 ± 31.4 | 670.1 ± 21.1 | 678.6 ± 17.3 | 670.5 ± 9.0 |
| Ct.Ar ($\text{mm}^2$) | 0.99 ± 0.11 | 0.94 ± 0.06 | 0.99 ± 0.17 | 0.90 ± 0.07 |
| Tt.Ar ($\text{mm}^2$) | 3.75 ± 0.20 | 3.67 ± 0.36 | 3.62 ± 0.43 | 3.57 ± 0.16 |
| Ct.Ar/Tt.Ar (%) | 26.5 ± 2.1 | 25.7 ± 1.6 | 27.1 ± 1.5 | 25.1 ± 1.4 |
| Ct.Th ($\mu\text{m}$) | 126.0 ± 9.3 | 122.0 ± 4.3 | 125.2 ± 11.4 | 118.6 ± 6.2 |
| Diaphysis (cortical) |  |  |  |  |
| vBMD ($\text{mg}/\text{cm}^3$) | 841.5 ± 15.0 | 834.6 ± 8.3 | 841.0 ± 16.7 | 824.6 ± 14.7 |
| Ct.Ar ($\text{mm}^2$) | 1.12 ± 0.10 | 1.04 ± 0.08 | 1.00 ± 0.18 | 0.93 ± 0.05 |
| Tt.Ar ($\text{mm}^2$) | 2.62 ± 0.22 | 2.42 ± 0.35 | 2.33 ± 0.37 | 2.27 ± 0.11 |
| Ct.Ar/Tt.Ar (%) | 39.7 ± 2.3 | 41.5 ± 3.5 | 41.8 ± 0.6 | 40.5 ± 1.6 |
| Ct.Th ($\mu\text{m}$) | 191.0 ± 10.5 | 193.0 ± 9.2 | 194.1 ± 20.4 | 186.7 ± 9.7 |
| Mid-diaphysis (cortical) |  |  |  |  |
| I<sub>ML</sub> ($\text{mm}^4$) | 0.22 ± 0.04 | 0.19 ± 0.05 | 0.18 ± 0.06 | 0.17 ± 0.02 |
| I<sub>AP</sub> ($\text{mm}^4$) | 0.55 ± 0.09 | 0.48 ± 0.10 | 0.45 ± 0.15 | 0.39 ± 0.03 |
| Femur Length (mm) | 16.4 ± 0.3 | 16.1 ± 0.8 | 15.9 ± 0.3 | 15.9 ± 0.2 |

a: $p < 0.05$ HFD vs. CFD (main effect)

While cancellous bone microstructure was substantially altered by HFD in the distal femoral metaphysis, cortical bone geometry remained similar between HFD and CFD in the distal metaphysis and diaphysis. In the femur, neither HFD nor exercise had a significant effect on cortical vBMD, cortical area (Ct.Ar), total area (Tt.Ar), cortical area fraction (Ct.Ar/Tt.Ar), or cortical thickness (Ct.Th) in either the cortical bone around the metaphyseal VOI or in the diaphyseal VOI (Table 1). Similarly, in the mid-diaphyseal VOI, neither HFD nor exercise affected medial-lateral moment of inertia (I_ML_, p = 0.25 and p = 0.38, respectively) or anterior-posterior moment of inertia (I_AP_, p = 0.11 and p = 0.28, respectively). Overall femur length was also similar across both diet (p = 0.17) and exercise (p = 0.52) groups (Table 1).

### 3.4 Cortical Bone Remodeling (Tibia)

Dynamic indices of cortical bone remodeling from dynamic histomorphometry were similar in the HFD and CFD groups at both the endosteal and periosteal surfaces, with no significant differences in MS/BS, MAR, or BFR/BS (Table 2). Exercise, however, did significantly affect the extent of active remodeling bone surface, with 62% greater endosteal MS/BS compared to sedentary groups (p = 0.016), but exercise did not affect endosteal MAR (p = 0.74) or BFR/BS (p = 0.57). The periosteal surface had little labeling, and neither HFD nor exercise had a significant effect on periosteal remodeling (Table 2).

**Table 2:** Dynamic Indices of Cortical Bone Remodeling in the Tibial Diaphysis (mean ± SD)

| Trait | Control Fat Diet |  | High Fat Diet |  |
| --- | --- | --- | --- | --- |
|  | Sedentary | Exercise | Sedentary | Exercise |
| Endosteal |  |  |  |  |
| MS/BS (%) | 27.8 ± 3.6 | 43.1 ± 13.4<sup>c</sup> | 27.0 ± 12.7 | 39.8 ± 6.8<sup>c</sup> |
| MAR ($\mu\text{m}/\text{day}$) | 0.59 ± 0.25 | 0.34 ± 0.24 | 0.33 ± 0.41 | 0.49 ± 0.36 |
| BFR/BS ($\mu\text{m}^3/\mu\text{m}^2/\text{day}$) | 0.30 ± 0.12 | 0.18 ± 0.13 | 0.13 ± 0.16 | 0.23 ± 0.16 |
| Periosteal |  |  |  |  |
| MS/BS (%) | 4.3 ± 5.1 | 13.4 ± 15.1 | 4.4 ± 6.7 | 8.7 ± 15.2 |
| MAR ($\mu\text{m}/\text{day}$) | 0.12 ± 0.16 | 0.22 ± 0.26 | 0.10 ± 0.13 | 0.20 ± 0.26 |
| BFR/BS ($\mu\text{m}^3/\mu\text{m}^2/\text{day}$) | 0.02 ± 0.02 | 0.09 ± 0.12 | 0.01 ± 0.01 | 0.05 ± 0.10 |

c: $p < 0.05$ Exercise vs. Sedentary (main effect)

### 3.5 Whole Bone, Apparent, and Tissue Mechanical Properties (Femur)

HFD negatively affected whole bone mechanical properties in the femur measured by three-point bending (Figure 6, Table 3). Compared to the CFD group, the HFD group had 18% lower yield load (p = 0.039) and nearly lower ultimate load (14% lower, p = 0.058) and stiffness (18% lower, p = 0.055). After accounting for body mass (ANCOVA), none of the mechanical properties differed between HFD and CFD groups, except whole bone stiffness tended to be lower in HFD compared to CFD even after body mass adjustments (p = 0.085, Table 3). Femoral length was similar across diet and exercise groups (Table 1), but when whole bone mechanical properties were adjusted for femur length (ANCOVA), none of the mechanical properties differed between HFD and CFD groups, except yield load was nearly lower in HFD compared to CFD (p = 0.082, Table 3). Similarly, none of the estimated apparent-level material properties – yield stress, ultimate stress, and Young’s modulus – were significantly affected by HFD or exercise (Table 3). Cortical tissue material properties assessed with nanoindentation were also not significantly affected by HFD or exercise (Table 3). Both hardness and reduced modulus values were consistent across regions (p = 0.66 and p = 0.42, respectively).

**Figure 6:**
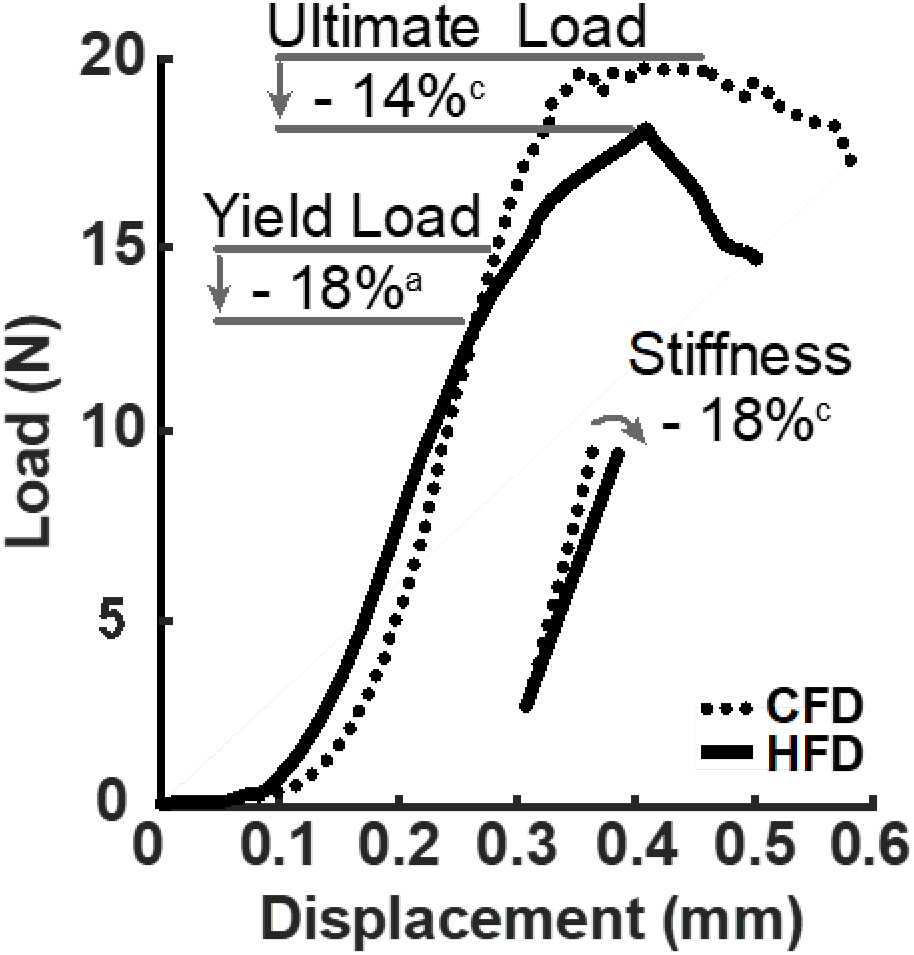
Representative force-displacement curves from femur three-point bending to failure. Relative to CFD, HFD significantly reduced yield load and nearly reduced stiffness and ultimate load. a: p < 0.05 HFD vs. CFD (main effect), d: p < 0.10 HFD vs. CFD (main effect).

**Table 3:** Whole Bone, Apparent, and Tissue Mechanical Properties in the Femoral Diaphysis.

| Trait | Control Fat Diet |  | High Fat Diet |  |
| --- | --- | --- | --- | --- |
|  | Sedentary | Exercise | Sedentary | Exercise |
| Whole bone mechanical properties (three-point bending) (mean ± standard deviation) | | | | |
| F<sub>yield</sub> (N) | 16.0 ± 2.1 | 16.1 ± 3.7 | 13.8 ± 2.3<sup>a,f</sup> | 12.6 ± 1.3<sup>a,f</sup> |
| F<sub>ult</sub> (N) | 22.3 ± 4.1 | 20.9 ± 2.1 | 19.1 ± 3.1<sup>d</sup> | 18.0 ± 1.6<sup>d</sup> |
| Stiffness (N/mm) | 137.5 ± 21.9 | 132.9 ± 17.6 | 119.3 ± 25.9<sup>d,e</sup> | 101.7 ± 26.2<sup>d,e</sup> |
| PYD (mm) | 0.13 ± 0.04 | 0.11 ± 0.05 | 0.11 ± 0.02 | 0.14 ± 0.02 |
| Work-to-fracture (mJ) | 6.09 ± 0.98 | 4.43 ± 0.97 | 3.79 ± 1.60 | 5.02 ± 0.75 |
| Apparent material properties (three-point bending) (mean ± standard deviation) | | | | |
| $\sigma_{\text{yield}}$ (MPa) | 93.4 ± 8.9 | 100.5 ± 22.4 | 96.5 ± 15.8 | 88.15 ± 5.0 |
| $\sigma_{\text{ult}}$ (MPa) | 117.6 ± 6.1 | 121.9 ± 11.2 | 117.2 ± 12.7 | 117.3 ± 6.9 |
| E (GPa) | 3.65 ± 0.15 | 4.15 ± 0.74 | 3.89 ± 0.73 | 3.58 ± 1.19 |
| Tissue material properties (nanoindentation) (least squares mean ± 95% confidence interval) | | | | |
| H (GPa) | 0.96 ± 0.21 | 0.85 ± 0.22 | 0.90 ± 0.20 | 0.87 ± 0.21 |
| E<sub>r</sub> (GPa) | 25.6 ± 5.6 | 22.7 ± 5.9 | 25.2 ± 5.6 | 22.6 ± 5.5 |

(main effects) a: $p < 0.05$ HFD vs. CFD raw data; d: $p < 0.10$ HFD vs. CFD raw data; e: $p < 0.10$ HFD vs. CFD body mass-adjusted; f: $p < 0.10$ HFD vs. CFD femur length-adjusted

### 3.6 Cortical Bone Composition (Femur)

HFD had only a small effect on tissue composition as assessed by Raman spectroscopy (Figure 7), nearly reducing carbonate substitution by 2% in the mid-cortex (p = 0.080) and by 3% along the periosteal edge (p = 0.083, Figure 7F). Exercise had more pronounced effects on cortical bone composition. Mineral maturity was nearly higher (2% greater phosphate crystallinity) for exercise groups compared to sedentary groups near the periosteal edge (p = 0.068, Figure 7A). Exercise did not affect mineral crystallinity in the mid-cortex (p = 0.81) or near the endosteal edge (p = 0.20). Mineral-to-matrix band intensity ratios near the endosteal edge were higher for exercise groups relative to sedentary groups for the phosphate/(proline+hydroxyproline) ratio (27% higher, p = 0.013, Figure 7B), phosphate/amide I ratio (18% higher, p = 0.030, Figure 7C), and phosphate/amide III ratio (25% higher, p = 0.023, Figure 7D). Similarly, the carbonate-to-matrix ratio (carbonate/amide I) near the endosteal edge was also increased for exercise compared to sedentary (13% higher, p = 0.023, Figure 7E). Carbonate substitution was not affected by exercise in any region (Figure 7F).

**Figure 7:**
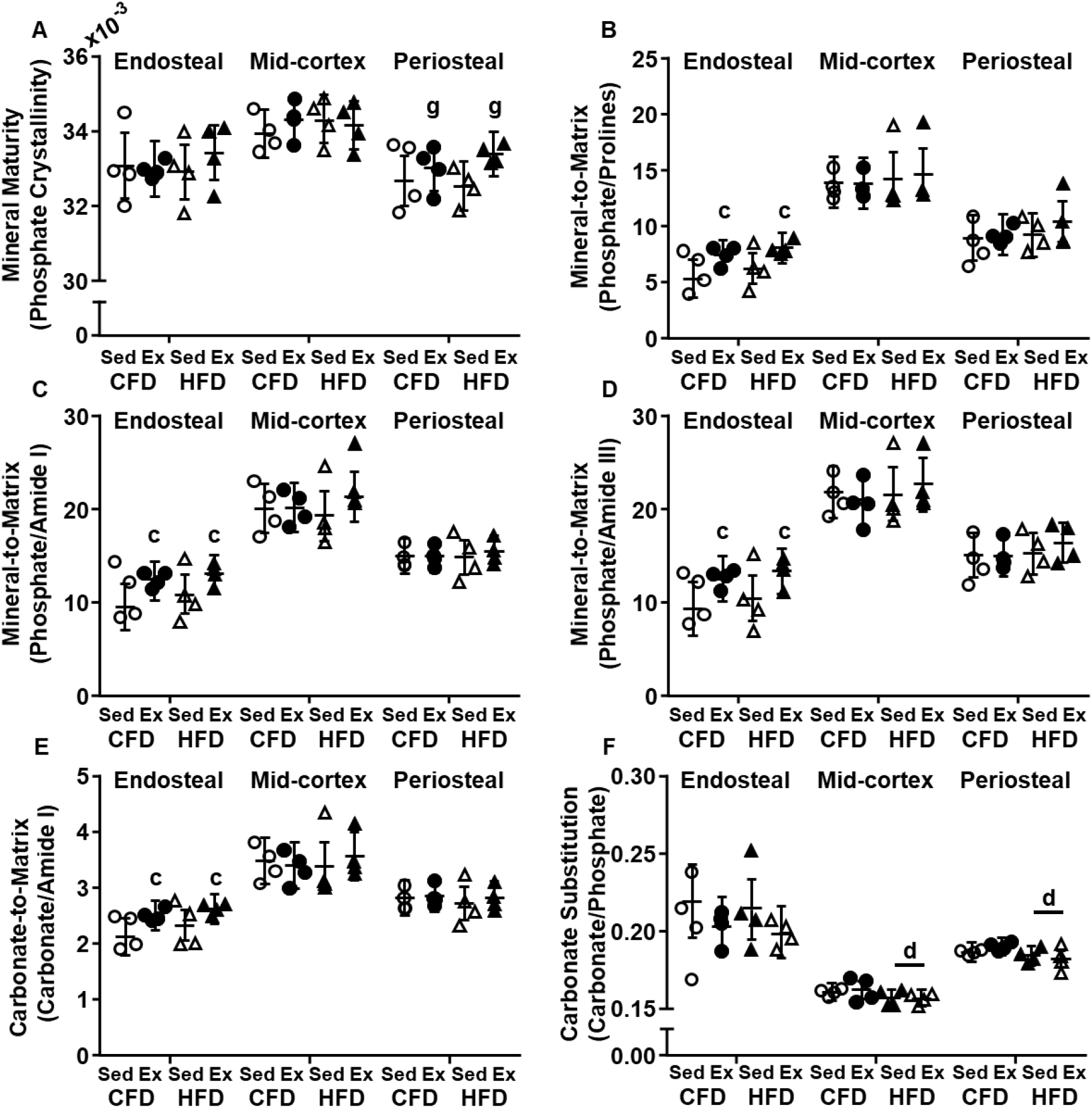
Relative to sedentary, exercise groups had increased A) mineral crystallinity on the periosteal edge. Along the endosteal edge, exercise increased B) phosphate *v1* to combined proline and hydroxyproline ratio, C) phosphate *v1* to amide I ratio, D) phosphate *v1* to amide III ratio, and E) carbonate *v1* to amide I ratio but not F) carbonate substitution. Points represent mean of all quadrants per femur, lines and bars represent estimated least-squares mean ± 95% confidence interval c: p < 0.05 Ex vs. Sed (main effect), d: p < 0.10 HFD vs CFD (main effect), g: p < 0.10 Ex vs. Sed (main effect).

### 3.7 Osteovascular Structure (Femur)

Osteovascular structure in the distal femoral metaphysis, as assessed by immunofluorescence, was not significantly affected by HFD or exercise (Table 4). Vessel area fraction (endomucin-positive blood vessels per total area) within the bone was similar HFD and CFD (p = 0.78) and between exercise and sedentary (p = 0.51) groups. Similarly, the average vessel-to-bone distance between endomucin-positive blood vessels and collagen type I-positive bone surfaces did not differ between HFD and CFD (p = 0.44) or between exercise and sedentary (p = 0.15). Bone surface area fraction (col-1-positive bone area per total area) was 32% lower in the HFD group compared to the CFD group (p = 0.034, Table 4), consistent with the reduced BV/TV noted above. Exercise did not affect col-1-positive bone surface area fraction (p = 0.51), also consistent with BV/TV results.

**Table 4.**
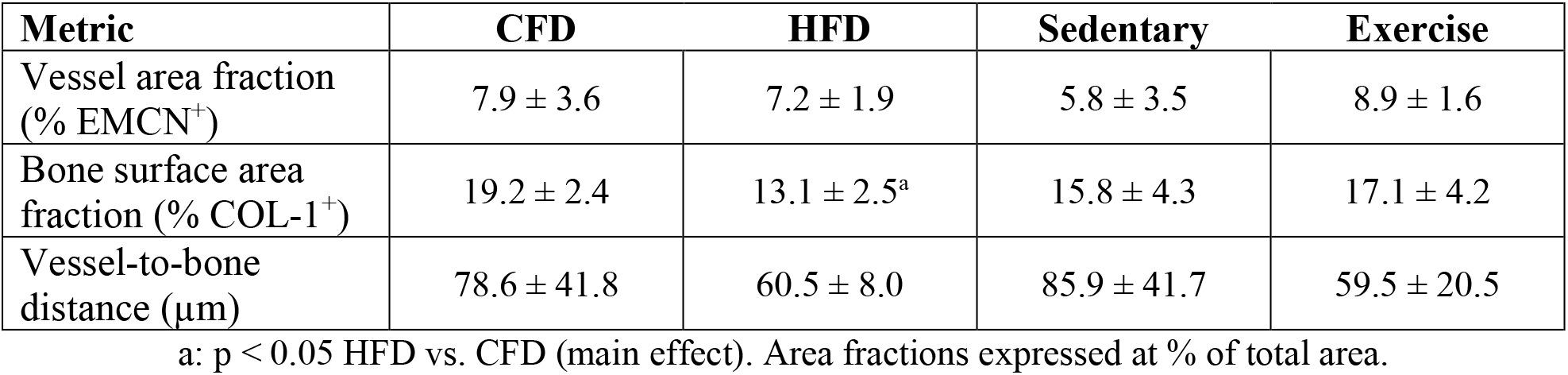
Osteovascular Structure in the Distal Femoral Metaphysis (mean ± standard deviation)

### 3.8 Osteovascular Crosstalk (Serum)

Crosstalk between endothelial cells and osteoblasts, as assessed by serum ELISA, was not affected by HFD or exercise (Table 5). At the end of the study, serum concentrations were similar between HFD and CFD groups (p = 0.27 for BMP2 and p = 0.89 for VEGF-A) and also between exercise and sedentary groups (p = 0.36 for BMP2 and p = 0.43 for VEGF-A).

**Table 5.** Serum Concentrations of Bone-Vascular Crosstalk Markers (mean ± standard deviation)

| Marker | Control Fat Diet |  | High Fat Diet |  |
| --- | --- | --- | --- | --- |
|  | Sedentary | Exercise | Sedentary | Exercise |
| BMP2 (pg/mL) | 80.7 ± 12.1 | 72.0 ± 7.5 | 73.8 ± 3.4 | 98.9 ± 31.0 |
| VEGF-A (pg/mL) | 42.8 ± 13.4 | 52.0 ± 8.8<sup>a</sup> | 55.4 ± 8.0 | 38.0 ± 8.5 |

## 4 Discussion

High fat diet-induced obesity reduced whole bone bending strength in the femur, without altering cortical bone mineral density, geometry, or apparent- or tissue-level material properties relative to control fat diet. Because bone strength depends on these parameters,^(9, 10)^ we expected one of them to be altered by HFD to explain the underlying cause for the relative strength deficits in that group. The reductions in bending properties with HFD were no longer significant after adjusting for body size, by including either body mass (yield and ultimate load) or femur length (ultimate load and stiffness) as a covariate. Although femur length was not significantly different between HFD and CFD groups, variations in body size seems to account for diet-related strength differences, as was also reported in a recent study where the magnitude of the effects of HFD on cortical area and bending strength were reduced after accounting for body mass.^(48)^ HFD had deleterious effects on cancellous bone microstructure in the distal femur, with reduced bone volume due to loss of trabeculae, which reduces bone strength to a much greater extent compared to trabecular thinning.^(49)^ Therefore, bone strength at primarily cancellous bone sites, like vertebrae, may also be reduced with HFD, as was demonstrated in the mouse L3 vertebra after HFD^(15)^ and the rat L6 vertebra after high sucrose diet induced-obesity.^(50)^ HFD did not alter osteovasculature in cancellous sites, with no differences in bone perfusion (proximal tibia) or vascular area and proximity to bone surfaces (distal femur) relative to CFD. This work reveals that HFD negatively affects cancellous bone microstructure without affecting vessel area, and cortical bone strength without affecting cortical geometry or material properties, and only slight changes to tissue composition.

HFD created an obese, hyperglycemic phenotype that persisted with daily treadmill exercise. After 9 weeks of diet, HFD groups were heavier than CFD groups, and after 13 weeks of diet, HFD groups had significantly lower glucose tolerance and weekly fasting serum glucose concentrations that were over 200 mg/dL, indicative of pre-diabetes.^(51)^ Although exercise had transient benefits to glucose tolerance in the HFD group, these benefits did not persist to the end of the study at Week 17, and daily treadmill exercise did not mitigate the negative effects of HFD on cancellous bone microstructure. Exercise had no effect on femoral cortical mechanical properties at the whole bone, apparent, or tissue levels, despite slightly increasing mineral-to-matrix ratios in the diaphysis. Exercise, but not HFD, increased the extent of active remodeling bone surface in the tibial diaphysis and bone perfusion in the proximal tibia but had no effect on the relative amount of blood vessels or the distance between blood vessels and bone surfaces in the distal femur.

High fat diet negatively affected cancellous, but not cortical, bone structure in the femur. Our reductions in cancellous microstructure and bone surface area in the distal femur with 60% fat diet from age 5-23 weeks are consistent with results from several other studies, which also reported cancellous bone degradation following HFD in young male C57Bl/6J mice. Compared to mice fed a control fat diet, mice fed a high fat diet (either 45% or 60% fat) from 3-6 weeks of age to 15-28 weeks of age experienced 18-49% reductions in cancellous bone volume fraction^(12,14,15,17,52)^ and 10-18% reductions in trabecular number^(13, 15)^ in the distal femoral metaphysis. Conversely, a 60% HFD from 7-28 weeks of age induced a 14% increase in trabecular cross-sectional area in the distal femur relative to CFD, but the measurements were obtained using peripheral quantitative computed tomography with a large voxel size (70 x 70 x 500 µm).^(53)^ Most studies have been performed in young, male mice, though a couple of studies found similar reductions in BV/TV in diets started after skeletal maturity was reached. A study comparing extended HFD from 7-28 weeks of age to short-term HFD from 25-28 weeks of age found a 19% decrease in cancellous BV/TV in the distal femoral metaphysis with extended HFD and a 12% decrease with short-term HFD.^(12)^ Similarly, a study comparing 60% HFD from 5-17 weeks of age (young) to HFD from 20-32 weeks of age (mature) found, compared to CFD mice of the same age, a 45% decrease in BV/TV in the distal femoral metaphysis in young mice and a 29% decrease in mature mice.^(15)^ These studies demonstrate that diet-induced obesity in male mice commonly leads to detrimental changes in cancellous bone microstructure, as we report here, and suggest that altered modeling during skeletal growth is not solely responsible for the negative HFD effects on microstructure.

The effects of HFD on cortical bone geometry in male C57Bl/6J mice are less consistent. Similar to our results, several groups report no effect on cortical bone parameters,^(13,14,16,54)^ but the study that reported increased trabecular cross-sectional area also found a 7% increase in cortical area in the diaphysis and a 21% increase in polar moment of inertia (pMOI) relative to CFD.^(53)^ Similarly, a 60% fat diet from 4-23 weeks of age resulted in an 11% increase in both diaphyseal Ct.Th and Ct.Ar relative to CFD,^(18)^ while a 60% fat diet from 6-18 weeks resulted in slightly expanded diaphyseal marrow area, lower Ct.Th, and similar pMOI relative to CFD.^(17)^ More research is required to determine specific underlying factors that may be contributing to this variability in HFD-induced effects on cortical bone structure, and whether these factors may help explain our reduced femoral strength. In particular, cortical porosity, which we could not examine at the resolution of our micro-CT scans, can impact bone strength,^(55, 56)^ and changes in cortical porosity with HFD are understudied. Two HFD studies have reported porosity measured with micro-CT using voxel sizes between 10-12 um,^(13, 57)^ but accurately measuring cortical porosity requires a higher resolution with a voxel size of 1-2 µm, particularly for small animals.^(58)^ To our knowledge only one study has examined porosity at this appropriate resolution, and they found that porosity measured with a 2-µm voxel size was up to 37% lower than porosity measured with a 1-µm voxel size, and that HFD reduced vascular canal porosity by 33% relative to CFD.^(59)^

HFD decreased whole bone mechanical properties in the femur, with 18% lower yield load, 14% lower ultimate load, and 18% lower stiffness in three-point bending compared to CFD. Other groups have also reported reduced femur bending properties for young male C57B/6J mice. Studies with HFD beginning at 3-6 weeks of age and ending at 19-28 weeks of age reported a 12% reduction in maximum load,^(19)^ 29% reduction in ultimate load, and 20% reduction in stiffness.^(52)^ Similar results have also been reported in the L3 vertebra, with mice fed a 60% HFD from 5-17 weeks of age (young) or from 20-32 weeks of age (mature) having 17-24% lower yield load, 16-26% lower maximum load, and 21-27% lower stiffness during compressive loading in both age groups compared to age-matched mice fed a CFD.^(15)^ Conversely, in a study of cantilever bending in the femoral neck, the HFD group (60% fat diet from 7-28 weeks of age) had 18% higher maximum load and 29% higher bending modulus compared to the CFD group.^(53)^

Despite reductions in whole bone mechanical properties, we found no changes in estimated apparent-level material properties with HFD. The study with reduced maximum force and stiffness in the L3 vertebra also found no significant changes in apparent-level material properties in either young or old HFD mice compared to age-matched CFD.^(15)^ Other groups have reported either reduced or increased apparent-level material properties for HFD vs. CFD in male C57B/6J mice. For whole femurs in three-point bending, two studies found that HFD (60% fat diet starting from 3-6 weeks-of-age to 19-28 weeks of age) caused 19-32% lower apparent elastic modulus, 15-26% lower maximum stress, and 24% lower yield stress,^(18, 19)^ while another study found 44% higher apparent elastic modulus.^(52)^ Tissue-level material properties were also unaltered by HFD in our study. To our knowledge, no previous study has examined the effects of HFD on tissue-level material properties. Since we did not find HFD-induced changes in bone density, structure, or tissue-level properties, the reduced whole bone strength may result from a combination of small changes in several parameters that were not statistically significant in this study.

Cortical tissue composition in the femur was altered by exercise, with increased mineral-to-matrix and carbonate-to-matrix ratios near the endosteal edge and increased mineral maturity near the periosteal edge. Mineralization of new bone tissue occurs slowly, so higher mineral-to-matrix and carbonate-to-matrix ratios are associated with older bone that is generally harder and stiffer.^(45, 60)^ However, a study using the same treadmill regimen initiated at 16 weeks of age found that treadmill exercise increased ultimate strain and the mineral-to-matrix ratio of phosphate *v*_1_ to summed proline and hydroxyproline without affecting tibial morphology, suggesting increased mineral-to-matrix ratios could be a mechanism by which bone adapts to exercise to maintain local functional strain.^(61)^ Other studies have used Raman spectroscopy to analyze the increased accumulation of advanced glycation end-products (AGEs), known to cause material differences that increase fracture risk,^(52,62–64)^ in rodent diabetic bone. Elevated glucose may lead to AGE accumulation in collagen,^(62, 65)^ which has been shown to increase resistance to plastic deformation and stiffness at the material level in bone.^(64, 66)^ A recent study in HFD mice (60% fat from 8-30 weeks of age) found no difference in mineral-to-matrix ratio, crystallinity, or carbonate substitution compared to CFD, but an increased amount of the AGE pentosidine (PEN), which was positively correlated with higher bending modulus despite lower stiffness and ultimate load.^(52)^ However, the Raman spectra in our study did not contain any of these AGE bands, indicating AGEs were not significantly present.

This study found no effect of HFD or exercise on 2D osteovascular structure (vessel area and proximity to bone surfaces) in the distal femur, but stereological methods are not ideal for measuring complex three-dimensional structures like the branching network of blood vessels,^(67)^ so HFD may have affected osteovascular parameters that are not quantifiable with stereology. Similar to our results, a recent study reported no HFD-related changes in the 3D vessel network in the proximal tibia using a new contrast agent with micro-CT.^(68)^ Specifically, HFD from 8-30 weeks of age did not affect the vessel volume per medullary volume or the distance between blood vessels and bone surfaces compared to CFD. However, this study also reported that HFD reduced the number of blood vessels by 3.9-fold and increased average vessel diameter by 2.7-fold, metrics that cannot be accurately quantified with stereological techniques.

Perfusion is a functional measure of blood supply to tissue that incorporates not only the amount of blood vessels but also the velocity and direction of the blood flow in the vessels, as well as vessel permeability and diameter.^(69)^ For example, if HFD increased vessel diameter but reduced vessel number compared to CFD, these changes could offset each other and result in the same perfusion measurement. Similarly, the increased perfusion observed with exercise may result from other structural changes to the vascular network besides vessel area and proximity to bone surfaces, which were similar between sedentary and exercise groups. Furthermore, bone perfusion likely experiences temporal changes in response to interventions like HFD and exercise; however, it was only measured at the end of the study to avoid causing inflammation in the hindlimb, as recommended by the group that developed the method for assessing perfusion in mouse tibiae.^(34)^

Diet-induced obesity has far-reaching physiological effects that can impact bone health and may be responsible for the observed envelope-specific changes to cancellous but not cortical bone structure. In this study, HFD led to the development of obesity and pre-diabetic levels of elevated serum glucose, both of which impact metabolic pathways that influence bone metabolism. Elevated glucose concentrations are associated with reduced BMD in rats and humans,^(70, 71)^ as well as *in vitro* proliferation and mineralization of osteoblasts.^(65,72,73)^ Obesity in humans and HFD-induced obesity mouse models are associated with increases in both leptin and glucocorticoids, which differentially affect cortical and cancellous bone envelopes.^(74–77)^ Leptin, which signals satiety, also promotes the maintenance of bone mass; when the leptin receptor is knocked out globally in mice they become obese, even without HFD, and gain cortical bone but lose cancellous bone.^(77)^ Mice with conditional knockout of the leptin receptor in bone marrow stromal cells, however, do not become obese without HFD. With 12 weeks of HFD, the conditional knockout prevented detrimental cancellous microstructure changes and decreased the number of mesenchymal stem cells (MSC) that differentiated into adipocytes compared to wild-type mice, suggesting obesity affects bone maintenance directly through leptin.^(78)^ Corticosterone, a glucocorticoid in rodents, is associated with increased bone resorption, but in growing mice the effect is bone- and site-specific, tending to increase endosteal resorption while preventing periosteal remodeling and leading to an expanded marrow cavity.^(76)^ Unlike leptin, the effect of obesity on increased serum glucocorticoids in either rodents or humans is unclear.^(75, 79)^ Lastly, increased amounts of marrow adipose tissue (MAT) may negatively affect cancellous bone structure. We did not quantify MAT in this study, but other groups report dramatic increases in the amount of metaphyseal MAT with HFD,^(13,24–27)^ and decreased MAT with intense exercise.^(24, 25)^ Moderate treadmill exercise did not affect bone microstructure in this study, but other studies that utilize more intense exercise regimen, such as free access to running wheel^(24,25,53,80)^ or high intensity treadmill training,^(81)^ found effects of exercise in HFD mice.

In conclusion, our study demonstrated that high fat diet-induced obesity caused detriments to cancellous bone microstructure and whole bone bending strength in the femur that were not concomitant with changes to metaphyseal perfusion or vascularity, or to cortical geometry or tissue properties. We also showed that moderate treadmill activity did not reverse the deleterious effects of HFD, increase intraosseous vascularity, or increase mechanical properties in this model. Exercise did, however, increase intraosseous perfusion in the tibia, and stimulate changes to tissue composition in the femur, without affecting geometry. These findings should be examined further by characterizing changes to intraosseous perfusion at different timepoints during the development of HFD, and by incorporating more intense exercise routines.

## Acknowledgments

We thank the following individuals for expertise and contributions to data collection: Eric Livingston and Dr. Ted Bateman (micro-CT); Keith Jones, Daniel Chester, and Dr. Ashley Brown (atomic force microscopy); Sandra Horton and Dr. Denis Marcellin-Little (histology); Sara Chopra, Nicholas Rinz, Dr. Roberto Garcia, Dr. Chuanzhen Elaine Zhou, and Dr. Fred Stevie (sample preparation, Raman spectroscopy); Dr. Eva Johannes (confocal fluorescence microscopy).

Research was supported by the Eunice Kennedy Shriver National Institute of Child Health and Human Development (NICHD) of the NIH under award number K12HD073945. The content is solely the responsibility of the authors and does not necessarily represent the official views of the NIH. This work was performed in part at the Analytical Instrumentation Facility (AIF) at North Carolina State University, which is supported by the State of North Carolina and the National Science Foundation (award number ECCS-1542015). The AIF is a member of the North Carolina Research Triangle Nanotechnology Network (RTNN), a site in the National Nanotechnology Coordinated Infrastructure (NNCI). The authors acknowledge the use of the Cellular and Molecular Imaging Facility (CMIF) at North Carolina State University, which is supported by the State of North Carolina and the National Science Foundation.

Authors’ roles: Study design: NH, AS, and JHC. Animal work: NH, AS, and EE. Data collection: NH, AS, JMC, EE, HT, MS, and SV. Data analysis: NH, AS, JMC, MS, and HT. Data interpretation: NH, AS, and JHC. Drafting and revising manuscript: NH, AS, JMC, and JHC. Approving final version of manuscript: NH, AS, JMC, EE, HT, MS, SV, and JHC. JHC takes responsibility for the integrity of the data analysis.

